# Quantifying the role of pre-existing tissue resident cellular immunity in limiting respiratory virus transmission

**DOI:** 10.1101/2025.09.14.676166

**Authors:** Ananya Saha, Sarah E Michalets, Ida Uddbäck, Hasan Ahmed, Jacob E Kohlmeier, Rustom Antia, Katia Koelle

## Abstract

Viral transmission from infected donors to uninfected recipients is the key event underlying the spread of viral pathogens at the level of a host population. Successful viral transmission from a donor to a recipient depends on several factors including the infectiousness of the donor. Donor infectiousness in turn can depend on the viral kinetics and viral load of the donor, donor behavior and symptoms, and donor immunity. Here, we use a mouse model of murine respirovirus (otherwise known as Sendai virus SeV) infection to quantitatively explore donor determinants of respiratory virus transmission. The experimental transmission studies we analyze are specifically designed to address the effect that pre-existing donor immunity may have on transmission potential by studying SeV transmission from both immunized and control (placebo-immunized) donors to naïve recipients. We specifically focus on the impact of tissue resident memory (TRM) CD8 T cells on donor transmission potential by considering immunization strategies that primarily generate CD8 T cell immunity. Through quantitative analyses of these experiments, we find that pre-existing CD8 TRMs act to reduce donor transmission potential. This reduction can be in part explained by a reduction in total infection load in immunized donors. However, even once differences in infection load between immunized and control donors are accounted for, immunized donors still have reduced infectiousness relative to control donors. We explore possible reasons for this unexpected pattern using a mathematical model that integrates within-host viral dynamics and between-host transmission occurrences. Analysis of model simulations, along with observations from knock-out experiments, suggests that interferon gamma (IFN-*γ*) may be partly responsible for the observed differences in infectiousness between control and pre-immune donors. Future experimental transmission studies should consider measuring IFN-*γ* levels when interpreting transmission outcomes in the context of host immunity.

## Introduction

Substantial transmission heterogeneity, or superspreading, underlies the dynamics of many infectious pathogens, such that the majority of secondary infections often stem from a small proportion of infected individuals [1, 2]. Transmission heterogeneity is particularly well documented in respiratory viruses such as SARS-CoV-2 [3, 4, 5, 6, 7, 8]. Understanding the sources of this heterogeneity is key to the design of effective intervention strategies. Many factors are known to contribute to transmission heterogeneity, including differences in host contact rates and in host infectiousness. Differences in host infectiousness are driven by differences in within-host viral kinetics as well as differences in disease symptoms such as coughing that can facilitate transmission [9, 10]. In turn, host factors such as host genetics, comorbidities, and pre-existing immunity impact an individual’s viral kinetics and symptom development and thus indirectly impact host infectiousness [11]. Here, we explore how pre-existing immunity of infected individuals may impact the transmission of a respiratory virus.

The role of pre-existing immunity in impacting virus spread has been increasingly studied over the last decade [12, 13, 14, 15]. This immunity can stem from either natural infection or vaccination. Studies focused on respiratory viruses have shown that immunity can reduce an individual’s susceptibility to infection [16, 17]. In the case of breakthrough infection, pre-existing immunity can reduce viral load, resulting in lower transmission [18]. Experimental transmission studies using animal models offer a tightly controlled approach for quantifying the role of pre-existing immunity in impacting transmission outcomes [12, 19] and dissecting the mechanisms of this impact. These studies provide an opportunity to track with high resolution the within-host viral dynamics of infected individuals that differ in their exposure histories and thus in their immunity. They also allow for quantitative assessment of how immunity impacts between-host transmission. Many different animal models have been used to study the transmission of respiratory viruses in experimental settings [12, 20, 21, 22, 23, 24]. A subset of these have specifically examined the effect of pre-existing immunity in reducing viral transmission [12, 19, 25, 26]. However, the majority of these studies have focused on antibody-mediated immunity rather than cellular immunity or have not been able to tease apart these two types of immunity in their impact on transmission.

Here, we specifically focus on quantifying the impact of pre-existing cellular immunity in donor individuals on their onward transmission potential of murine respirovirus using analyses of two different experimental transmission studies performed in the same laboratory. Murine respirovirus (Sendai virus, or SeV) is a respiratory virus of mice that, like influenza A viruses and coronaviruses, is both airborne- and contact-transmitted in mice. Both studies characterize onward SeV transmission from mice that are intranasally vaccinated with a T-cell vaccine (hereafter, “pre-immune” mice) versus from placebo-vaccinated (hereafter “control”) mice to naïve contact mice. Both of these studies benefit from *in vivo* imaging of viral infection in live mice to longitudinally quantify within-host infection burden. By mapping the relationship between within-host infection burden and transmission probability, our analyses first indicate that pre-existing cellular immunity reduces infection burden and therewith reduces the probability of onward transmission. More intriguingly, our analyses also indicate that cellular immunity results in lower infectiousness than expected once reduced infection burden has been accounted for. We interpret these findings in the context of a mathematical model that considers within-host viral and immune dynamics to shed light on the mechanisms by which pre-existing T-cell immunity may impact onward transmission potential.

## Results

### Pre-existing cellular immunity reduces onward transmission probability by reducing infection burden

In the first experimental transmission study, infected index mice were placed in cages with naïve contact mice for a fixed time window of two days, with exposure to infected index mice occurring during different time windows following donor infection. Index mice comprised two different groups: a pre-immune group that had been vaccinated with a Sendai Virus (SeV) T-cell vaccine and a control group that had been given a placebo vaccine that did not elicit a T-cell response to SeV. The experimental design is summarized in the *Materials & Methods* section and schematically shown in Figure S1. We hereafter refer to this study as the “Short & Fixed” study because contact mice were caged with index mice for short, fixed time periods of two days.

Figure 1A shows infection dynamics of the control and the pre-immune index mice using *in vivo* measurements of bioluminescence flux, which can be interpreted as a measure of total infected cell numbers in an infected mouse. In the control mice, flux increased until 3-5 days post infection and then declined after day 5. By day 7, flux returned to levels that indicated resolution of infection. When the control index mice were caged with naïve contact mice during the 1-3 days-post infection time window (D1-3), transmission efficiencies were 3/4, 4/4, and 1/4 in the three replicates performed (Figure S2A). When these same control index mice were caged with naïve contact mice during the D3-5 window, transmission efficiencies were 4/4 in each of the three replicates (Figure S2A). Finally, during the D5-7 window, transmission efficiencies were 2/4, 1/4, and 3/4 in the three replicates (Figure S2A). Because infection levels are higher during the D3-5 window than in either the D1-3 window or the D5-7 window (Figure 1A), these results indicate that infection levels in a donor may be positively related to transmission probability. To assess this possibility more quantitatively, we first calculated the infection burden for each of the control index mice for each of the three time windows (see *Materials & Methods*). We then plotted each contact’s infection outcome (0 for no infection; 1 for infection) against the infection burden of its corresponding index animal over the time period of its exposure (Figure 1B). A logistic regression model fit to these data indicates that transmission probability from an index animal is positively associated with its infection burden (Figure 1B) (p-value = 0.013).

**Figure 1:**
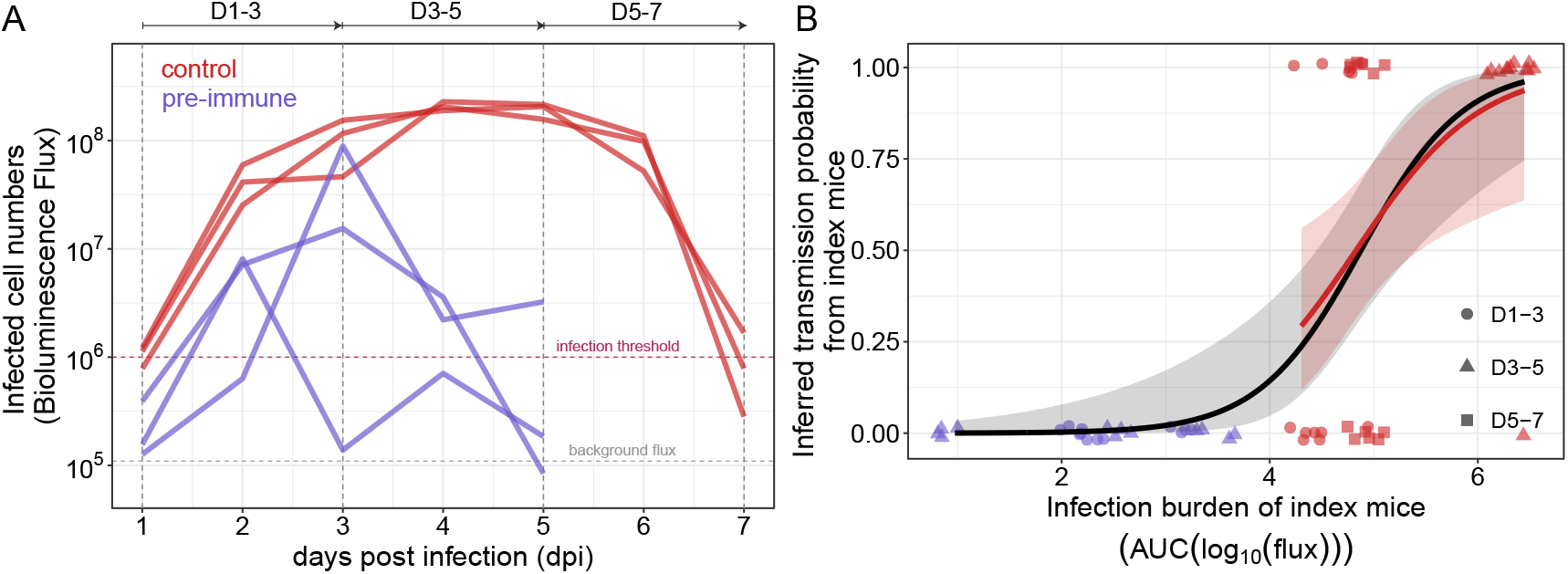
Infection dynamics and transmission outcomes from control and pre-immune index mice in the “Short & Fixed” study. **(A)** Infection dynamics of control index mice (red) and pre-immune index mice (purple). Infection levels are measured using bioluminescence flux (photons/sec) and plotted as a function of days post index mouse infection. Vertical dashed lines and the arrows on top of the figure panel specify the time windows when the index mice were co-housed with the naïve contact mice. **(B)** Transmission outcomes and estimated transmission probabilities as a function of index animal infection burden. Data points show transmission outcomes of individual contact mice (0 for no infection; 1 for infection), with their color reflecting whether the index mouse was from the control group (red) or from the pre-immune group (purple) and their marker reflecting the time window of exposure. Logistic regression using all of data points is shown in black. The grey shaded regions show the 95% confidence intervals in the estimated transmission probabilities. Logistic regression using only the control data points is shown in red. On the log odds scale, the estimated parameters for the black curve are - intercept: -9.94 (SE: 2.93), slope: 2.04 (SE: 0.61); and the estimated parameters for the red curve are - intercept: -8.08 (SE: 3.33), slope: 1.67 (SE: 0.67).

We repeated this analysis for the three pre-immune index mice. In these mice, flux again increased following infection, but peak flux occurred earlier (2-3 days post infection) and was generally lower than in the control index mice. Infection in pre-immune animals also resolved earlier (by day 5) than in the control index mice. None of the three pre-immune index animals transmitted SeV infection to any of the contact animals in either the D1-3 window or the D3-5 window (Figure S2B). A D5-7 window was not evaluated, given resolution of infection of the pre-immune index animals by day 5. We calculated infection burdens for the pre-immune index animals during the D1-3 windows and the D3-5 windows, and plotted the contacts’ infection outcomes (all 0) against the infection burdens of their corresponding index over the exposure time period (Figure 1B). Given the lack of transmission, we could not fit a logistic regression to these data alone. However, we could combine the transmission outcomes from the control index mice and the pre-immune index mice and fit a logistic regression to this combined dataset. Doing so, we again found a significant positive association of transmission probability with total infection burden (Figure 1B) (p-value = 0.0008).

It is apparent from Figure 1B that pre-existing T-cell immunity reduces infection burden, which in turn results in a reduction of transmission probability. As such, the lower transmission efficiencies from pre-immune index mice relative to those from control index mice appears to be, at least in part, driven by a reduced infection burden in the pre-immune mice.

### Pre-existing cellular immunity reduces infectiousness in addition to infection burden

Because the “Short & Fixed” study resulted in no transmission from the pre-immune index mice, we next considered a second study where the duration of exposure to both pre-immune index mice and control index mice was extended, thereby increasing probabilities of onward transmission. Previously analyzed in [27], this second study considered three groups of index mice: a placebo-vaccinated control group (as in the “Short & Fixed” study) and two different pre-immune groups. One pre-immune group was vaccinated intranasally with the same T-cell SeV vaccine that was used in the “Short & Fixed” study. The second pre-immune group was vaccinated intraparetonially with this vaccine. The experimental design of this study is further summarized in the *Materials & Methods* section and schematically shown in Figure S3. Here, to parallel the index groups in the “Short & Fixed” study, we limited our analyses of this transmission study to the control group and the intranasally vaccinated pre-immune group.

In this second study, infected index mice were transferred into cages with naïve contact mice either at 1 day post-infection (dpi) (D1+ group), at 3 dpi (D3+ group), at 5 dpi (D5+ group), or at 7 dpi (D7+ group). The index mice were then kept in their cages for the remainder of the experiment (up to 14 dpi). We hereafter refer to this study as the “Long & Variable” study because contact mice were caged with index mice for variable time periods spanning 7 to 13 days (longer than the 2 days in the “Short & Fixed” study). Figure 2A shows infection dynamics of the control and the pre-immune index mice again using measurements of bioluminescence flux. As was observed in Figure 1A, flux in the control index mice increased until 4-5 days post infection and then started declining after day 5. Flux returned to levels that indicated resolution of infection by day 7-9. SeV transmission from the control index mice to naïve contact mice occurred during the D1+, D3+, and D5+ exposure windows with high transmission efficiency (Figure S4A). No onward transmission occurred during the D7+ exposure window (Figure S4A).

**Figure 2:**
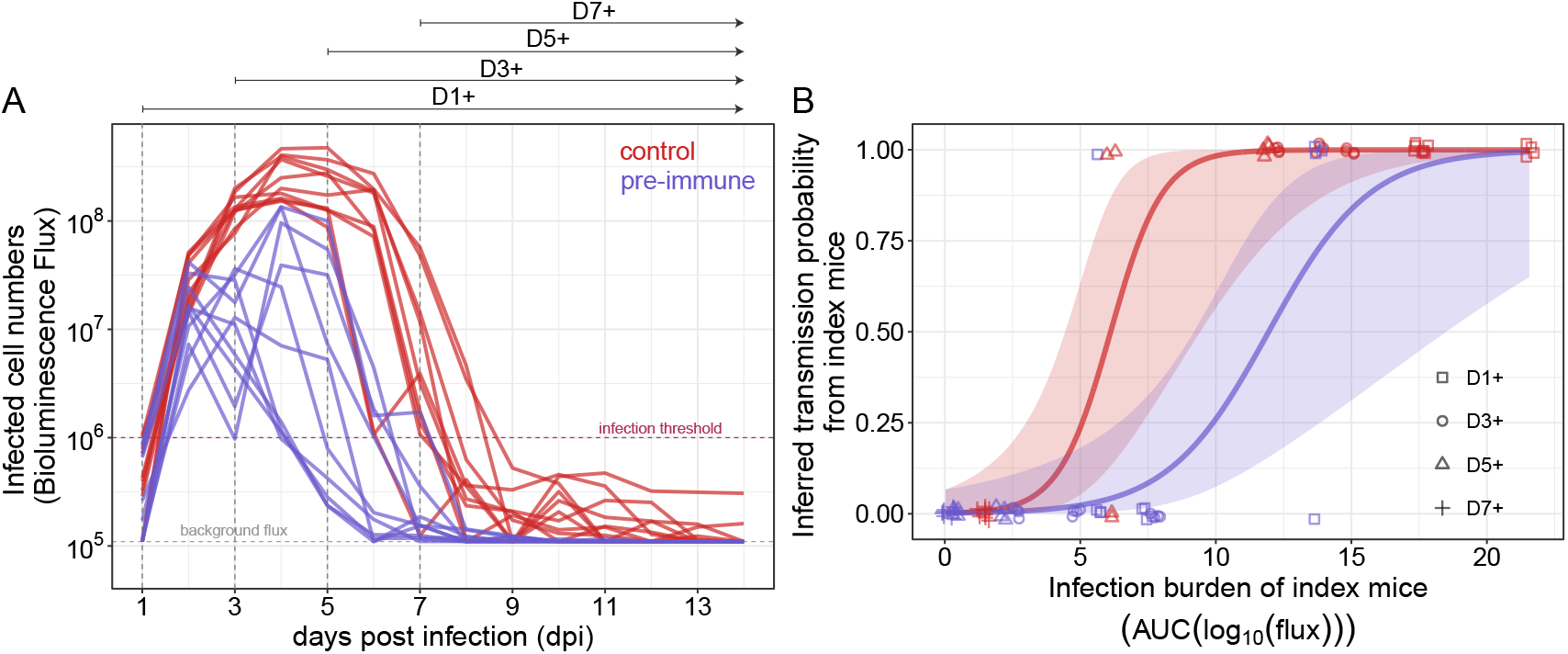
Infection dynamics and transmission outcomes from control and pre-immune index mice in the “Long & Variable” study. **(A)** Infection dynamics of control and pre-immune index mice. Infection levels were measured using bioluminescence flux and plotted as a function of days post infection. Vertical dashed lines and the arrows on top of the figure panel specify the time windows when the index mice were co-housed with the naïve contact mice. **(B)** Transmission outcomes and estimated transmission probabilities as a function of index animal infection burden. Data points show transmission outcomes of individual contact mice (0 for no infection; 1 for infection), with their color reflecting whether their corresponding index mouse was from the control group (red) or from the pre-immune group (purple) and their marker reflecting the time window of exposure. A logistic regression model was fitted to all the data points with immune status as a covariate to estimate the functional relationship between transmission probability and infection burden. On the log odds scale, the estimated parameters for the red curve are - intercept: -6.4 (SE: 1.9), slope: 1.04 (SE: 0.34); and the estimated parameters for the purple curve are - intercept: -6.4 (SE: 1.9), slope: 0.53 (SE: 0.18). Both relationships are statistically significant: p-value = 0.002 for the control group and p-value = 0.003 for the pre-immune group.

Consistent with the infection dynamics in the “Short & Fixed” study, flux in the pre-immune index mice peaked earlier (2-4 days post infection) and at levels that were lower than in the control group (Figure 2A). Infection in pre-immune animals again resolved earlier (by day 5-7). Figure S4B shows the infection dynamics of the pre-immune contact mice alongside the infection dynamics of their corresponding index mice. Some transmission from pre-immune index mice to contact mice occurred in the D1+ exposure window, but no transmission in D3+ and D5+ exposure windows was observed.

To quantify the relationship between infection burdens in the index mice and their onward transmission probabilities for this transmission study, we first calculated the infection burden of the index mice during their respective transmission windows. We then plotted each contact’s infection outcome against the infection burden of its corresponding index animal for both control and pre-immune groups (Figure 2B). Logistic regression models fit to these data again indicate that transmission probabilities are positively associated with infection burden. However the extent of this association is different between the control and the pre-immune groups (Figure 2B). Specifically, at intermediate index mouse infection burdens, it appears that the probability of transmission is lower from pre-immune index mice than control index mice for the same infection burden.

In a final analysis, we combined the available data from the “Short & Fixed” and the “Long & Variable” studies to ascertain the robustness of our results and further quantify the extent to which pre-immunity might reduce donor infectiousness. We again fit a logistic regression to infer the relationship between donor infection burden and transmission probability, this time with the combined data points (Figure 3A). Consistent with our previous results shown in Figure 2B, we find that the transmission probability from pre-immune index animals is estimated to be lower than that of control index animals when controlling for infection burden.

**Figure 3:**
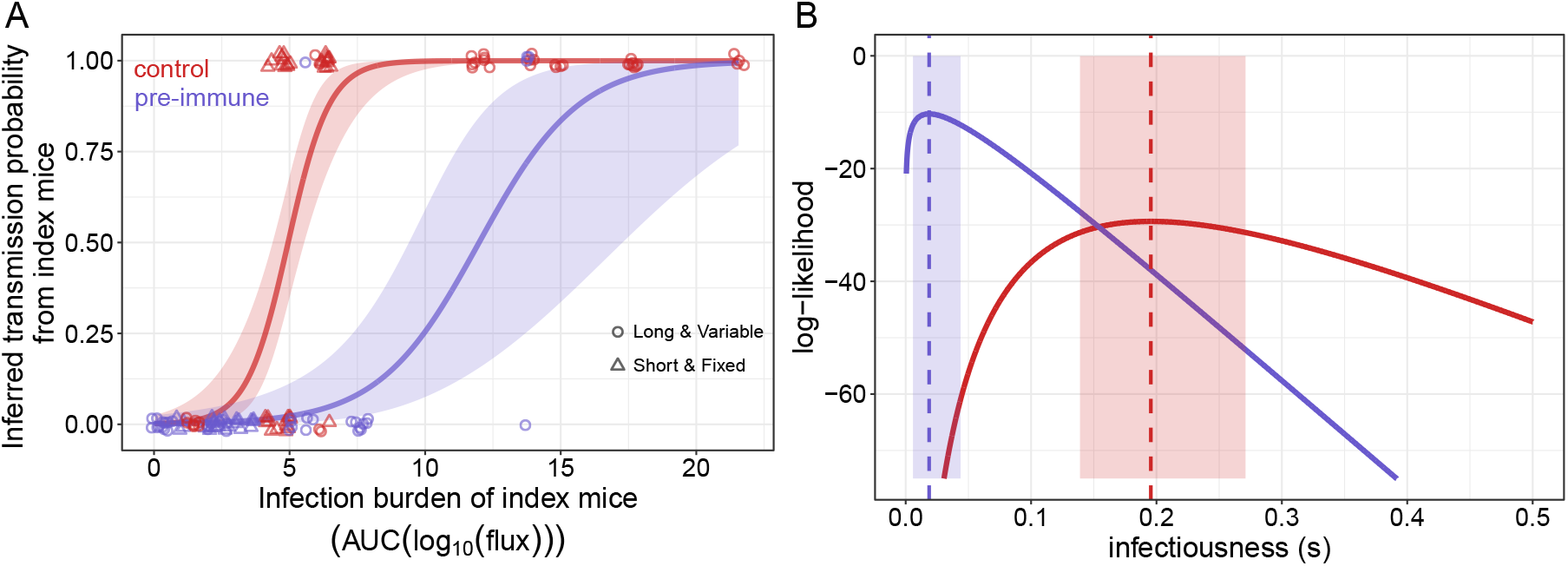
Transmission outcomes from and infectiousness of control and pre-immune index mice when combining data from the “Short & Fixed” and “Long & Variable” studies. **(A)** Transmission outcomes and estimated transmission probabilities as a function of index animal infection burden. Data points show transmission outcomes of individual contact mice (0 for no infection; 1 for infection), with their color reflecting whether the index mouse was from the control group (red) or from the pre-immune group (purple) and their marker reflecting the study from which the data points are derived. A logistic regression model is fitted to all the data points with index immune status as a covariate. On the log odds scale, the estimated parameters for the red curve are - intercept: -6.46 (SE: 1.4), slope: 1.33 (SE: 0.28); and the estimated parameters for the purple curve are - intercept: -6.46 (SE: 1.4), slope: 0.51 (SE: 0.14). Both relationships are statistically significant: p-value = 1 × 10^−6^ for the control group and p-value = 3 × 10^−4^ for the pre-immune group. **(B)** Estimates of infectiousness *B* of the control index animals (red) and the pre-immune index animals (purple). Red and purple dashed vertical lines show maximum likelihood estimates of the parameter *B* for the control group (*B*_*control*_) and the pre-immune mice (*B*_*pre*–*immune*_), respectively. Maximum likelihood estimates are given by the values of *B* that correspond to the peak values of the log-likelihood curves shown. 95% confidence intervals are shown as shaded regions surrounding the maximum likelihood estimates of *s*.

These results indicate that infection burden is unlikely to be the sole predictor of transmission probability. Indeed, previous analyses of several experimental transmission studies have posited that infection stage, the onset of clinical manifestations, the virus’s ability to successfully establish a new infection, and the onset of the immune response might be additional factors that impact transmission probabilities [28, 29, 30, 31]. We therefore further quantitatively explored the extent to which pre-existing TRMs reduce host infectiousness using a time-varying force-of-infection (FOI) inference approach (see *Materials & Methods*). Here we considered a simple time-varying FOI approach that uses longitudinal measurements of infection burden in an index animal and infection outcome of its corresponding contact animals to statistically estimate a parameter *s* that quantifies index infectiousness. Application of this approach to the data from the two transmission studies indicates that the infectiousness of a pre-immune index mouse is lower than the infectiousness of a control index mouse (Figure 3B).

We can biologically interpret this finding as follows: a pre-immune index mouse is expected to have a lower probability of transmission to a contact mouse than would a control index mouse that has the same measured infection burden. Moreover, conditional on successful transmission, transmission from a pre-immune index mouse would be expected to occur later than from a control index mouse that has the same infection dynamics. Next we use experimental observations along with mathematical modeling to explore possible reasons of the inferred differences in infectiousness.

### IFN-*γ* appears partly responsible for the inferred differences in infectiousness between control and pre-immune groups

One possible reason for lower inferred infectiousness of the pre-immune group could be related to the production of functional virus that can transmit from index mice to successfully infect contact mice. Production of virus from infected cells might be different at different time points following infection due to the onset of host immune responses or the presence of pre-existing resident memory T cells. To evaluate this possibility, we therefore sought to correlate functional viral load, measured by plaque assay, to infected cell numbers, measured by bioluminescence flux, in the control and the pre-immune groups. We did not find any evidence that the flux-PFU relationship was different between when memory T cells were present relative to when they were absent (Figure S5). This suggests that pre-existing tissue resident memory T cells reduce virus replication and facilitate early clearance of the virus, which results in reduced infection burden of the pre-immune host. This reduced infection burden is reflected in both flux (i.e., infected cell numbers) and pfu (i.e., functional viral load) measurements. After accounting for this reduced infection burden, intriguingly, we still infer an additional reduction in infectiousness from the pre-immune group.

A second possible reason for the inferred differences in infectiousness could be the rapid production of IFN-*γ* by resident memory T cells following infection. Presence of IFN-*γ* in the pre-immune group might render functional virus to be less infectious, for example, by transmitting IFN-*γ* when transmitting the virus. Here we quantitatively explore this possibility using mathematical modeling. We first consider a within-host mechanistic model of acute infection and then predict between-host transmission probabilities with the modeled infection dynamics. The mathematical description of the within-host model is provided in *Materials & Methods*. Briefly, uninfected target epithelial cells become infected with Sendai virus following exposure to free infectious virus. With a time-delay of ∼1 day, infected cells release type-I interferon (IFN). Recognition of infected cells further stimulates existing tissue resident memory CD8 T cells (TRM). In turn, stimulated TRM cells produce effector molecules such as perforin that directly kill infected cells. With a time-delay on the order of hours, interactions between infected cells and T cells (when present) result in IFN-*γ* production from stimulated T cells [32]. The overall IFN response in turn blocks virus production from infected cells. This model captures the essential mechanistic details following an acute viral infection, both in the absence and in the presence of existing TRMs. The model also reproduces key features of infected cell dynamics that we observe using bioluminescence flux in both the control and pre-immune index mice (Figures 4A, 4B). Parameterized for pre-immune index mice, this model predicts peak infected cell numbers that are about one order of magnitude smaller than the peak in control index mice. Model simulations further predict that the peak in pre-immune mice occurs approximately a day earlier than that of control mice and that infected cell clearance also occurs earlier in pre-immune mice. Figure S6 shows the dynamics of the other model variables. Furthermore, by combining our simulated flux dynamics with corresponding infectiousness profiles (estimated in Figure 3), we could reproduce observed patterns of onward transmission probabilities in both the “Short & Fixed” and “Long & Variable” studies (Figure S7). For instance, in Figure 4C, we show that our transmission probability predictions for the D3+ group from the “Long & Variable” study reproduce observed transmission probabilities: with control mice, we predict close to a 100% transmission probability, whereas with the pre-immune mice, largely because *s*_*pre*–*immune*_ *< s*_*control*_, we predict only a ∼12% transmission probability. Overall, these results support our data analyses in Figure 3 and suggest that pre-existing TRMs reduced transmission probability to an extent that cannot be solely explained by reductions in infection burden.

**Figure 4:**
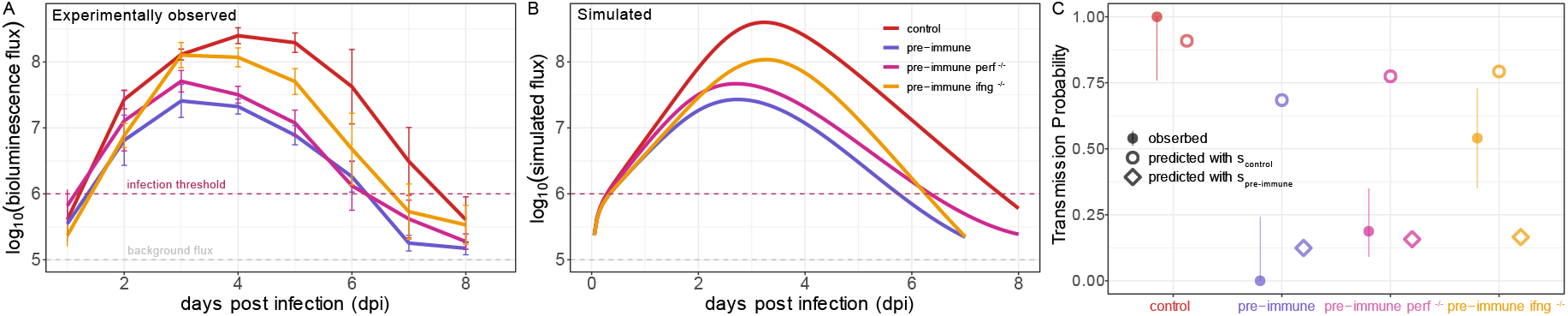
Within-host infection dynamics and transmission probability estimates under different experimental conditions. (**A**) Experimentally observed infection dynamics under different conditions are plotted in different colors. Color key is provided in panel B. (**B**) Simulated infection dynamics under these different conditions generated by changing parameters in our within-host model. (**C**) Observed (solid circles with error bars) and model-predicted (open markers) transmission probabilities for the **D3+** group from the “Long & Variable” transmission study under different experimental conditions. Perf^−/–^ was simulated by reducing the model parameter *σ*_*M*_. IFN-*γ*^−/–^ was simulated by setting the T cell IFN-*γ* production rate to 0 in the within-host model. Transmission probabilities were estimated for the different pre-immune groups using both *B*_*control*_ (open circles) and *B*_*pre*–*immune*_ (open diamonds) to examine what causes reduced infectiousness in the pre-immune group.

We then used our model to quantitatively explore different mechanisms of protection conferred by tissue resident memory T cells in limiting transmission. Using the within-host model, we first simulated a case where perforin was knocked-out from pre-immune mice (pre-immune perf^−/–^). Since perforin is not the only cytotoxic effector molecule by which CD8 T cells exhibit their cytolytic functions, we implemented perf^−/–^ by reducing the T-cell killing rate (*σ*_*M*_) 3 fold in the model equations. Simulations of this model yields infected cell number dynamics that are similar to those simulated for the no knock-out pre-immune group (Figure 4B), with only slightly higher infected cell numbers throughout the course of simulated infection. These model simulations further tightly recover observed infection dynamics under perf^−/–^ experimental conditions, shown in Figure 4A. Under this knock-out scenario, using the estimated *s*_*pre*–*immune*_ from Figure 3B, we see only a slight increase in the predicted transmission probability from the pre-immune group compared to what is observed under the no knock-out condition (purple vs magenta diamonds). This result is in line with transmission probabilities observed in the experiments by Uddbäck and colleagues [27], where they tracked transmission from perf^−/–^ pre-immune mice for the D3+ group in the “Long & Variable” transmission setting (purple vs magenta solid circles with error bars). As such, the lower infectiousness of pre-immune mice cannot be parsimoniously explained by the impact that tissue resident memory T cells have via perforin.

We next simulated a case where IFN-*γ* was knocked-out from the pre-immune mice (pre-immune IFN-*γ*^−/–^). Since we assumed that IFN-*γ* is produced only by TRM cells when present, this knock-out was simulated by setting the T-cell IFN-*γ* production rate (*y*) to 0 in the model equations (see *Methods*). With IFN-*γ*^−/–^, we see a more substantive increase in simulated infected cell numbers compared to the perf^−/–^ simulations (Figure 4B). We further see a slightly delayed peak relative to simulations parameterized for the no knock-out pre-immune group (Figure 4B). Our model’s predictions of higher infected cell numbers and a slightly delayed peak are reproduced under IFN-*γ*^−/–^ knock-out experimental conditions (Figure 4A). We then again estimated transmission probabilities under this knockout scenario by combining model simulations with infectiousness estimates from Figure 3B.

Using the estimated *s*_*pre*–*immune*_ from Figure 3B, we again predicted only a slightly increased transmission probability under perf^−/–^ conditions in the pre-immune group compared to no knock-out conditions in the pre-immune group. This prediction, however, does not match the findings by Uddbäcket and colleagues that reported a significantly increased transmission probability from the D3+ group with IFN-*γ* knock-out (Figure 4C). Underestimation of the observed transmission probability in the pre-immune IFN-*γ*^−/–^ case could be due to IFN-*γ* being a factor governing infectiousness in the pre-immune group. In the absence of IFN-*γ*, infectiousness is increased in this group. By using the estimated *s*_*control*_ from Figure 3B for the pre-immune IFN-*γ*^−/–^ group, our prediction of transmission probability is only slightly higher than the observed transmission probability (open and solid golden yellow circles with error bars), suggesting that IFN-*γ* could be largely responsible for the observed differences in infectiousness between the control and pre-immune groups.

## Discussion

Transmission of pathogens from infected individuals to uninfected hosts causes spread of infectious diseases in populations of individuals. In this study, we explored the role of immunity in preventing or limiting transmission of respiratory viruses. We used a combination of statistical data analyses and mathematical modeling to dissect the role of tissue-resident memory T cells (TRM) in limiting transmission of infectious virus between hosts. Specifically, we analyzed experimental data of Sendai virus (SeV) transmission in a mouse model in the presence and absence of TRMs.

Our earlier analysis of this transmission system suggested that with pre-existing TRMs, SeV transmission is curtailed earlier than without any T cell immunity [27]. Here we wanted to quantify this effect of T-cell immunity by specifically linking within-host infection dynamics to between-host transmission. Towards that end, we conducted further transmission experiments using a modified design of “Short & Fixed” exposure duration (Figure 1) and quantified the relationship between transmission probability and total within-host infection burden in this modified setting. Contrasting results from our previous transmission setup with “Long & Variable” exposure duration (Figure 2) [27] to those from our modified setup, we observed that tissue resident memory T cells differentially modulated the relationship between transmission probability and total within-host infection burden when exposure duration was extended and variable (Figure 2). Further investigations of this differential effect mediated by TRMs suggested that pre-existing TRMs also reduce infectiousness per unit infection burden (Figure 3).

Our different infectiousness estimates for the control and the pre-immune groups were based on relating transmission probabilities to the log-transformation of the total bioluminescence flux. It needs to be determined in future studies if other functional forms also provide similar results. The choice of functional form to link within-host infection dynamics to between-host transmission is an active research area and more mechanistic details about early infection processes will be required to derive an accurate mathematical description.

Using mathematical modeling, we further explored possible within-host immunological mechanisms that could be responsible for the apparent reduced infectiousness of pre-immune mice. Earlier studies have developed models to study acute infection dynamics, mainly to understand what processes are responsible for clearing infections [33, 34]. Other studies have used models to estimate infection-related parameters that are difficult to quantify experimentally [35]. The model developed in [35] was extended in later studies to further consider immune responses and other complexities not captured by the earlier model [36, 37, 38]. A recent study has specifically modeled the role of CD8 T cells in clearing influenza virus infection in mice and linked within-host viral dynamics to lung injury and disease severity [39]. Another study specifically modeled SeV infection dynamics in mice that also reported bioluminescence flux and PFU titers to reflect infected cell and virus dynamics [40]. This study modeled the death rate of infected cells as a density-dependent term to reflect a biphasic decline of infection. Here we used a model structure similar to that reported in [35] (see *Methods* for detailed model description). Our rational to choose this much simpler model is because (i) we do not see any biphasic infection clearance as observed in [40]; (ii) we wanted to model the effect mediated by respiratory track resident TRMs which act mainly by producing IFN-*γ* [27, 32]; (iii) in the absence of dynamical data on specific immune responses, we wanted to adopt the most parsimonious approach to capture the essential features of the within-host infection dynamics. The model was parameterized to reflect the experimentally observed infection dynamics.

Further, the purpose of the modeling and simulation was to capture patterns seen in the data collected at different scales, that is, both within hosts and transmission between hosts. Previously, only a few studies have attempted to quantitatively link within-host infection dynamics to transmission outcomes [29, 30, 31, 41, 42]. Here, we linked within-host infection dynamics to transmission probabilities for SeV in mice using a force of infection (FOI) framework. We could broadly recapitulate experimentally observed transmission dynamics (Figure S7). Here we would like to note that our modeling framework was not developed to match the actual numbers of different quantities that were measured experimentally. Transmission probability estimates in the experiments are not exactly the same as we predicted, and could be simply due to nonrandom contact patterns in a cage and other transmission-related heterogeneity [43, 44]. Further, when predicting transmission probability, we used only a single set of within-host model parameters along with single values for infectiousness (either *s*_*control*_ or *s*_*pre*–*immune*_). Almost certainly there will be variability in these parameters, which will impact the predicted mean transmission probability.

Acute infections are hypothesized to be controlled and eventually cleared primarily by innate immunity, such as interferon, NK cells, and recruitment of other inflammatory cytokines, in the absence of an adaptive immune memory [45, 46]. Adaptive immune memory, i.e., antibodies or memory CD8 T cells, may further facilitate early clearance of infection and limit transmission [27, 47, 48, 49, 50]. Earlier studies have looked into the role of pre-existing antibody responses in reducing SeV reinfection in mice, following their exposure to SeV by different transmission modes [51, 52]. In our system, there was no pre-existing SeV specific memory B cells present. This allowed us to specifically look at the role of memory CD8 T cell response. Previously a study by Price and colleagues investigated the role of CD8 T cells in limiting transmission of influenza viruses [53]. They found that a vaccine, inducing a T-cell response, could limit virus replication in the nasal cavity compared to unvaccinated animals, whereas virus dynamics in the lung was similar between these groups. Importantly, they observed significantly reduced transmission from the T-cell vaccine group. T cells play crucial roles in clearance of virus infections by directly killing infected cells and by producing antiviral cytokines. Moreover, an interferon response is stimulated by virus-infected cells and pre-existing T cells can bolster this effect furthermore. Zhou et al. [54] tested the role of IFN-*γ* to blocking infection in the presence of SeV specific CD8 T cells generated by Ad-SenNP vaccination. They found that CD8 T cells mediate early resistance to viral challenge primarily through release of IFN-*γ*. Our findings of reduced infectiousness (and essentially no transmission at early time points following challenge infection) along with reduced infection burden in the pre-immune group could also be due to an early IFN-*γ* response that might reduce the infectious viral load even if virus is detected using a plaque assay. It is poorly understood how functional virus detected by plaque assay relate to infectiousness of an individual leading to effective between host transmission [11, 28]. We are currently unsure of the exact biological mechanisms of how IFN-*γ* can modify pre-immune donor infectiousness and future studies should address this question in more detail.

T cells provide important antiviral defense and also target evolutionary conserved epitopes on viral genomes. This makes them an excellent candidate for incorporating into the design of universal vaccines against rapidly evolving respiratory viruses. The role of T cell mediated protection against respiratory viruses is understudied compared to the role of antibodies. Our earlier study along with this study focused on understanding the role of T cells in limiting transmission and reducing susceptibility to infection. However, we have used mouse and murine para influenza virus as our model system. Whether similar findings can be obtained in other animal models and other respiratory viruses needs to be determined. Future studies investigating the role of T cells in other animal models that can closely reflect respiratory virus transmission in humans are therefore needed to assess whether and to what extent T cell immunity can protect against infection and onward transmission potential.

## Materials and Methods

### Experimental transmission study details

The “Short & Fixed” transmission study and the previously published “Long & Variable” transmission study were both conducted in the Kohlmeier research laboratory at Emory University. Details on the “Long & Variable” transmission study are already provided in [27]. The “Short & Fixed” transmission study also used six- to eight-week-old female C57BL/6 mice obtained from Jackson Laboratory and housed them under the same pathogen free conditions at Emory University. Both transmission study experiments were completed in accordance with the Institutional Animal Care and Use Committee guidelines of Emory University, PROTO201700581. Age matched mice were randomly assigned to experimental groups for both experiments.

For intranasal priming, 30 plaque-forming units (PFU) Influenza A/Puerto Rico/8/34 (PR8-WT), 50 PFU Influenza A/Puerto Rico/8/34 expressing Sendai nucleoprotein FAPGNY-PAL epitope (PR8-SenNP) were administered in a 30 *μl* volume under isoflurane anaesthesia (Patterson Veterinary). Encoding luciferase SeV (Sendai-Luc) was generated and grown as previously described [27]. For direct Sendai-Luc infection, 1500 PFU in 30 *μl* was administered intranasally under isoflurane anaesthesia.

*In vivo* imaging was done using an In Vivo Imaging System (IVIS) Lumina LT Series III (Perkin Elmer) with an XFOV-24 lens as previously described [27]. Bioluminescent signal was quantified by manually drawing regions of interest around the respiratory tract using known anatomical markers. Bioluminescence in this system quantifies the number of infected cells, rather than extracellular virus, as it relies on reporter gene expression and is only detectable when the viral genome is translated.

For longitudinal detection of viral titers from the nasal cavity of infected mice, virus was collected by dipping the nose of each mouse into 1% BSA in PBS 20 times in a 12-well plate under isoflurane anaesthesia as described previously [27]. Samples were collected in duplicates. Plaques were counted using the following formula: PFU/ml = (average number of plaques) × (dilution fold).

### Transmission study design

We designed a transmission experiment setup where SeV transmission to naïve contact mice from an immune or a naïve index mouse is tracked for a short and fixed contact duration distributed throughout its entire infection period. Throughout this paper, we refer to this experiment as the “Short & Fixed” duration transmission experiment (experimental design shown in Figure S1). As depicted in the schematic, we first vaccinated mice intranasally with PR8 WT or with PR8 SendNP recombinant vaccine. Vaccination with PR8 WT generated no SeV specific T cell immunity, whereas the PR8 SendNP i.n. vaccination generated Sendai NP specific resident memory CD8 T (TRM) cells [27]. 35 days following vaccination, mice were artificially infected with SeV. Each of the infected index mice was then placed in cages with four naïve contact mice for a period of 2 days distributed throughout its entire infection period. After the end of each 2 day-period, the index mice were placed in new cages with a fresh and unexposed group of contact mice (Figure S1). Thus transmission from index mice was tracked within 1-3 days post infection (dpi) (D1-3 group), and then within 3-5 dpi (D3-5 group), and finally within 5-7 dpi (D5-7 group). Since the pre-immune index mice cleared the infection (defined by bioluminescence flux level below the infection threshold) at day 5, transmission was not tracked afterwards.

The second transmission study, referred to as “Long & Variable” duration transmission is schematically represented in Figure S3. The detailed description of this transmission experiment is provided in our previous work by Uddback and Michalets et al. [27].

### Data analysis and modelling of transmission experiments

#### Logistic regression, transmission probability, and infectiousness calculations

We first calculated the area under the curve (AUC) of the bioluminescent signal, measured by the IVIS, of an index mouse from the time when the mouse was caged with contact mice to the time the mouse was kept in the cage to track transmission. During the contact period between an index mouse and its corresponding contact mice in each cage, we then analyzed which of the 4 contact mice got infected using a threshold of infection *log*_10_(*flux*) = 6. The threshold of infection was estimated as the 95% CI of the background bioluminescent signal from two uninfected mice [27]. By tracking each contact mouse placed in a given cage with a given index mouse, we could then estimate the probability of transmission from that index mouse as a function of its AUC and immune status using a simple logistic regression model. Fit to the logistic regression was performed using the R function *glm()* with family as *binomial* and link as `logit.` More specifically, here, probability of transmission is calculated as a two-parameter logistic function with intercept *a* and slope *b* according to the formula

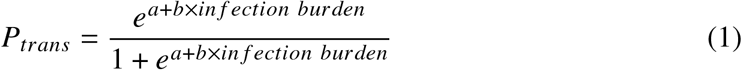

where infection burden is the AUC of the *log*_10_(*flux*) for the corresponding exposure duration.

For infectiousness calculations, assuming transmission from an index to a contact follows a poisson process,

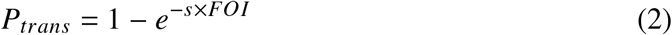

where *s* is infectiousness. We consider a simple definition of the FOI for a given transmission window *t*_1_ to *t*_2_ as 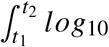 (*in fection load*) *dt*. This translates to taking the AUC of the *log*_10_ infected cell numbers (or fllux) of an index mouse for it’s given transmission window. In this formulation of modeling transmission as a poisson process, *s* is equivalent to *b* in the logistic model described above in equation 1. The parameter *s* is then estimated using maximum likelihood [55] by calculating the probabilities of the contact animals having been infected when they are caged with an index mouse across a range of *s* values. For all these calculations, the background AUC which is determined by the background flux of 10^5^ was subtracted from the calculated AUCs to infer the effect of actual infection burden.

Any contact mouse, that showed detected infection by crossing the threshold of infection more than 3 days after their corresponding index mouse was removed or cleared infection, was removed from analysis.

### Within host mathematical modeling

Here we use a simple mathematical modelling framework analogus to the target cell limitation model, borrowed from the literature [35]. The model equations are as follows,

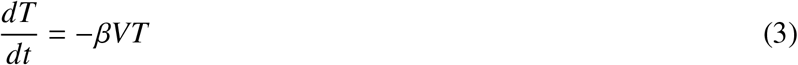

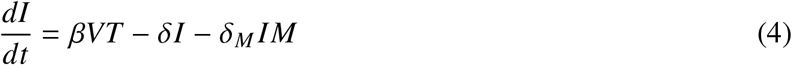

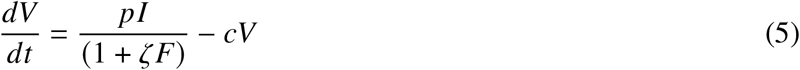

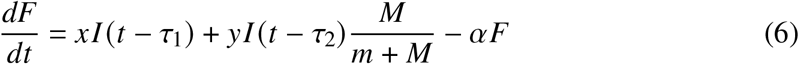

The target cell population *T* is infected by the virus *V* following a mass-action law with an infectivity of *β*. Upon infection of uninfected cells, infected cells can be cleared at a per capita rate of *δ*. Infected cells can also be killed by the TRM cells (*M*) at a second order rate constant *δ*_*M*_. Infected cells produce virus at a per capita rate of *p*. However, the production rate of virus from infected cells might be reduced in the presence of interferon, modelled here as *F*. The effect of interferon on the virus production rate is modelled using the term 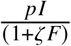, where *ζ* is efficacy of interferon in blocking virus production from the infected cells. Note, here we do not model *M* as a dynamical equation since previous study by McMaster et al., [32] reported that upon infection the TRM cell population doesn’t divide and proliferate, rather act by secreting IFN-*γ* rapidly. Finally, we model the type-I interferon (IFN) response to accumulate following a time delay of *τ*_1_ proportional to the infected cell population at (*t* – *τ*_1_), and IFN-*γ* to be produced by the TRM cells (M) following a time delay of *τ*_2_ when TRM cells are present. IFN-*γ* production is also proportional to the infected cell numbers at (*t* – *τ*_2_), but depends on the TRM cell numbers. Interferon (both type-I and IFN-*γ*) is cleared at a per capita rate of *α*. In this model, we have lumped the effects of type-I and IFN-*γ* into a single equation of *F*. It might be the case that these two interferon subtypes play different roles and show different kinetics. But in the absence of more granular data on interferon dynamics, we preferred this approach.

The infection related parameters to simulate this model were partly informed by Pinky et al. [40]. Note that we do not use the exact same model structure and the same parameter values as used in Pinky et al. [40] since we don’t see a biphasic clearance pattern of infection in the mouse strain that was used for our study. Different infection kinetics with differences in inoculum dose and mouse strain was reported earlier in Burke et al. [51]. Parameters related to IFN dynamics have been informed by work done by Handel and colleagues [56], since they modeled influenza infection in mice accounting for IFN dynamics. Specifically, we use the following parameter values: *β* = 1 × 10^−4^ (*virus*)^−1^ *day*^−1^, *p* = 2.8 (*cell*)^−1^ (*day*)^−1^, *c* = 4 *day*^−1^, *τ*_1_ = 1.5 *day, τ*_2_ = 0.2 *day, δ* = 1.7 *day*^−1^, *δ*_*M*_ = 0.003 *day*^−1^*cell*^−1^, *x* = 0.018, *y* = 0.005, *α* = 0.8 *day*^−1^, *ζ* = 2, *m* = 10 *cells*. The model was simulated with the following initial conditions: *T*_0_ = 10^5^ *cells, V*_0_ = 5, *I*_0_ = 0, *F*_0_ = 0, and *M* = 300 [27] if there is pre-existing TRM cells, otherwise *M* = 0.

Following initiation of an infection, simulations were stopped when the infected cell numbers went below 2 cells reflecting stochastic extinction and/or elimination of a small number of infected cells by the pre-existing TRM cells. After simulating the model with the above mentioned parameters, we calculated simulated flux using *log*_10_(*simulated flux*) = *log*_10_(*I*)+ *log*_10_(*background flux*). We then calculated the transmission probabilities for any exposure window by integrating the simulated flux within that window following the FOI framework using equation 2.

## Acknowledgments

Research reported in this publication was funded by the National Institute of Allergy and Infectious Diseases, Centers of Excellence for Influenza Research and Response, contract number 75N93021C0001.

