## Supplemental Figures S1 to S7 for "Quantifying the role of pre-existing tissue resident cellular immunity in limiting respiratory virus transmission"

### Supplementary materials

Figs. S1 to S7

All data and code used in this analysis will be posted online.

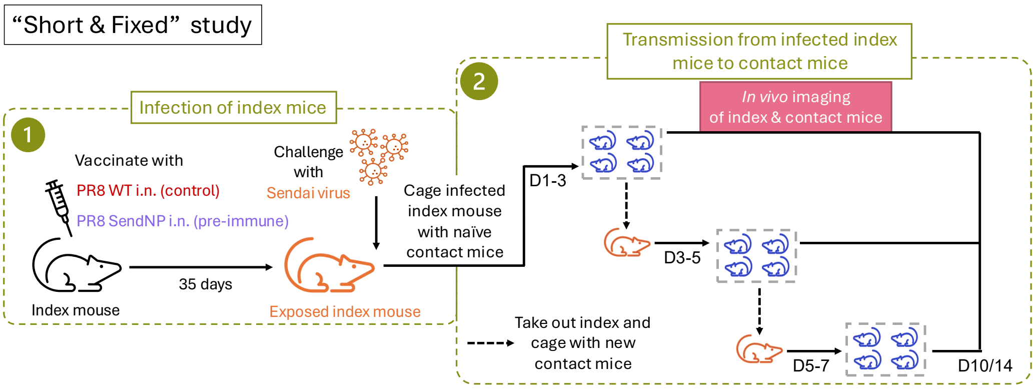


**Figure S1**: **Schematic of the "Short & Fixed" experimental transmission study.** Index mice were grouped into a control and a pre-immune group. Mice in the control group were intranasally (i.n.) inoculated with a placebo vaccine (PR8 WT). Mice in the pre-immune group were intranasally (i.n.) inoculated with a vaccine that elicited a cellular immune response to Sendai virus (PR8 SendNP). After 35 days, index mice from both the control group and the pre-immune group were experimentally challenged with Sendai virus. Each index mouse was then sequentially placed in three cages, each of which had 4 naïve contact mice. The sequential placement of index mice occurred on days 1-3 (D1-3), days 3-5 (D3-5), and days 5-7 (D5-7). Infection dynamics in both index and contact mice where measured daily using bioluminescence flux.

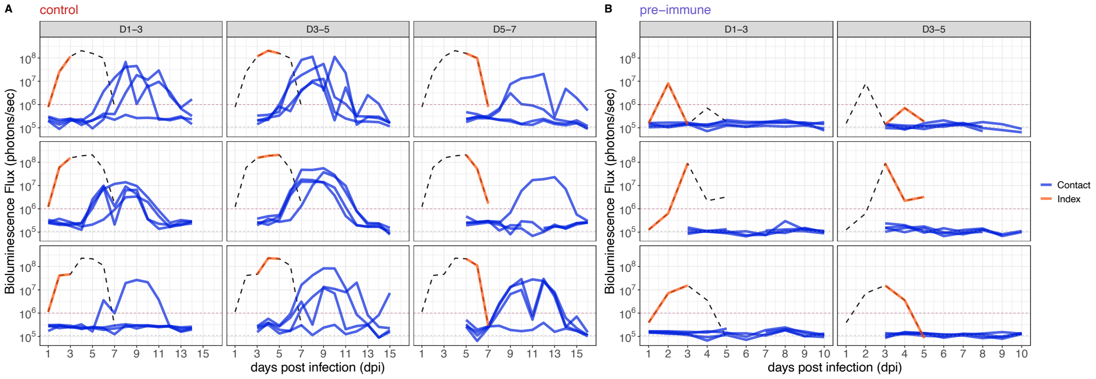


**Figure S2**: **Paired infection dynamics of infected index mice and their corresponding contacts in the "Short & Fixed" study. (A)** Paired infection dynamics of the control index mice and their contacts. **(B)** Paired infection dynamics of the pre-immune index mice and their contacts. The red horizontal dashed line in each panel shows the bioluminescence flux threshold for calling infection. The grey horizontal dashed line in each panel shows the background level of bioluminescence flux. Solid orange lines in the index mice infection dynamics represent their exposure duration in each transmission window.

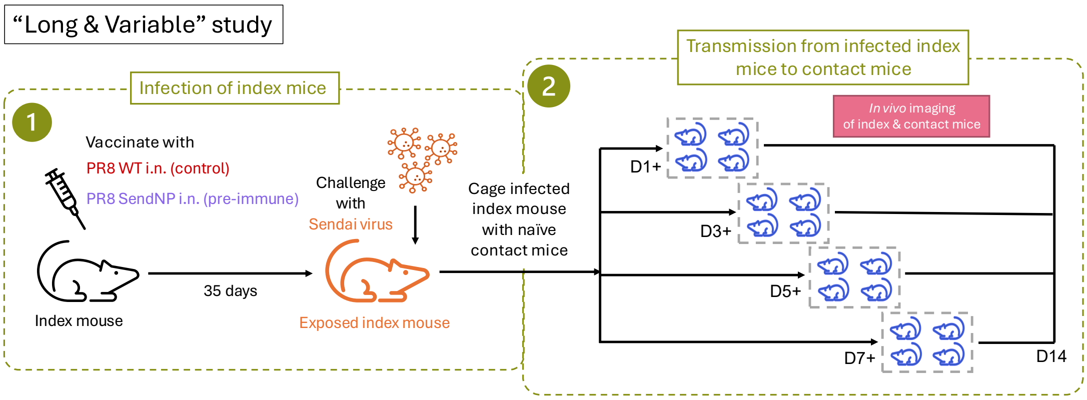


**Figure S3**: **Schematic of the “Long & Variable” experimental transmission study.** Index mice were grouped into a control and a pre-immune group. Mice in the control group were intranasally (i.n.) inoculated with a placebo vaccine (PR8 WT). Mice in the pre-immune group were intranasally (i.n.) inoculated with a vaccine that elicited a cellular immune response to Sendai virus (PR8 SendNP). After 35 days, index mice from both the control group and the pre-immune group were experimentally challenged with Sendai virus. Each index mouse was then placed in a cage with 4 naïve contact mice starting at either at 1 (D1+), 3 (D3+), 5 (D5+), or 7 (D7+) days post infection (dpi). Infection dynamics in both index and contact mice where measured daily using bioluminescence flux.

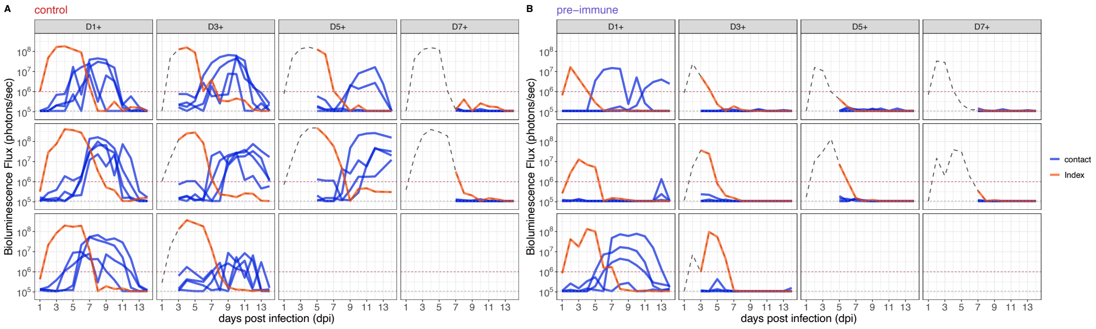


**Figure S4**: **Paired infection dynamics of infected index mice and their corresponding contacts in the “Long & Variable” study.** **(A)** Paired infection dynamics of the control index mice and their contacts. **(B)** Paired infection dynamics of the pre-immune index mice and their contacts. The red horizontal line in each panel shows the bioluminescence flux threshold for calling infection. The grey horizontal line in each panel shows the background level of bioluminescence flux. Solid orange lines in the index mice infection dynamics represent their exposure duration in each transmission window.

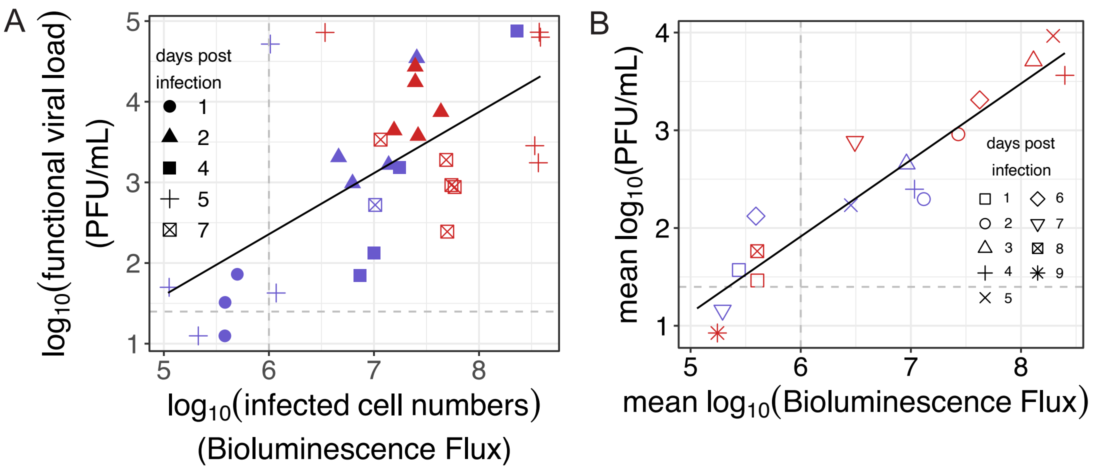


**Figure S5**: **Relationship between the number of infected cells (measured using bioluminescence flux) and the amount of infectious virus (measured using plaque assays).** We wanted to test if the relationship between the infected cell numbers and infectious virus titers is changed with and without an immune response. To test if this is the case, we performed nasal wash plaque assays from pre-immune and naive mice at different days post infection (dpi) with Sendai Virus. **A** We tested the relationship between the plaque forming units (a measure of functional virus) and the IVIS flux levels (a measure of infected cell numbers) from paired data where the nasal wash was collected and bioluminescence flux was measured from the same mouse at the same dpi. With pre-existing T cell memory, the bioluminescence flux levels are decreased and we see an equivalent decrease in the measured levels of plaque forming units. We would like to note here that the PFU measurements are noisy and the adjusted $R^2$ value for a straight line fit to this data is 0.39. **(B)** The data used for this analysis was taken from [27]. A group of vaccinated (PR8 SendNP i.n.) and unvaccinated (PR8 WT i.n.) mice were infected and daily measurements of flux using IVIS were performed. A separate group of vaccinated or unvaccinated mice were infected and daily measurements of PFU/mL were performed using the nasal wash plaque assay. Hence, here the measurements for flux and PFU/mL do not come from the same mouse. A linear relationship fitted between the mean $log_{10}(flux)$ and the mean $log_{10}(PFU/mL)$ is indicated by the black line.

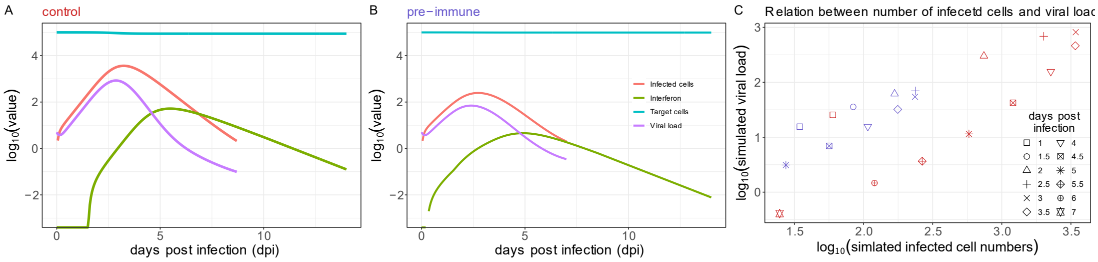


**Figure S6**: **Simulations of within-host viral and immune dynamics. (A)** Dynamics of target cells (blue), infected cells (red), viral load (magenta), and interferon (olive green) from the within-host model simulated with 'control' parameters. **(B)** Dynamics of target cells (blue), infected cells (red), viral load (magenta), and interferon (olive green) from the within-host model simulated with 'pre-immune' parameters. **(C)** The relationship between simulated infected cell numbers and viral load from the control simulation (A) and pre-immune simulation (B). In the pre-immune group, we see lower infecetd cell and correspondingly lower viral load compared to the control group. However, the two groups show similar patterns in relating viral load to total infected cell numbers.

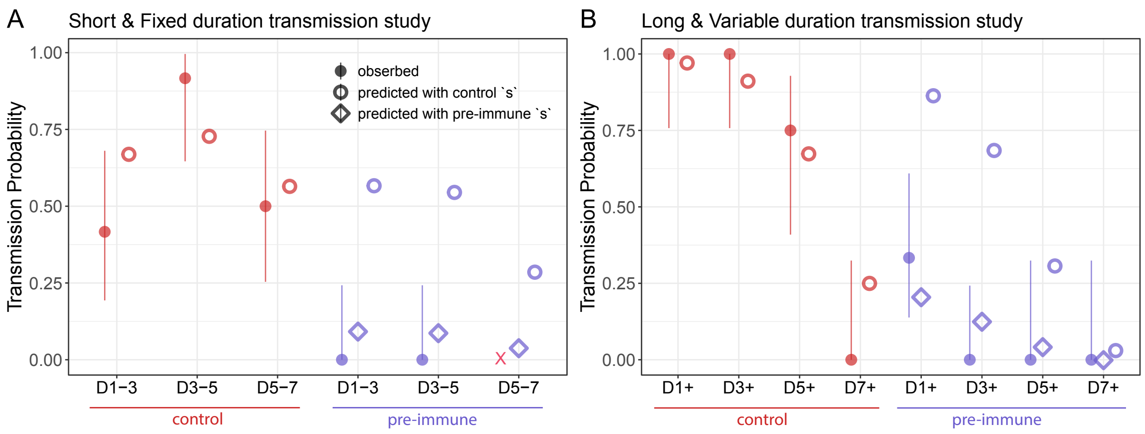


**Figure S7**: **Model prediction and validation with transmission experiment data.** **(A)** Comparison between observed and predicted transmission probabilities for the "Short & Fixed" study and **(B)** "Long & Variable" study. Experimental data was not available for the D5-7 transmission window for the pre-immune group. For the pre-immune group, we used both $s_{control}$ (in open circles) and $s_{pre-immune}$ (in open diamonds) to predict transmission probability with simulated pre-immune infection dynamics. This allowed us to study the effect of TRMs on the infection burden alone, keeping infectiousness the same as the control group. As seen in the plots, for both transmission experiments, a reduction in infectiousness for the pre-immune group was necessary to predict the observed transmission probabilities.
